# Accurate non-invasive quantification of astaxanthin content using hyperspectral images and machine learning

**DOI:** 10.1101/2024.09.23.614444

**Authors:** Marco L. Calderini, Salli Pääkkönen, Aliisa Yli-Tuomola, Hemanta Timilsina, Katja Pulkkinen, Ilkka Pölönen, Pauliina Salmi

## Abstract

Commercial cultivation of the microalgae *Haematococcus pluvialis* to produce natural astaxanthin has gained significant traction due to the high antioxidant capacity of this pigment and its application in foods, feed, cosmetics and nutraceuticals. However, monitoring of astaxanthin content in cultures remains challenging and relies on invasive, time consuming and expensive approaches. In this study, we employed reflectance hyperspectral imaging (HSI) of *H. pluvialis* suspensions within the visible spectrum, combined with a 1-dimensional convolutional neural network (CNN) to predict the astaxanthin content (μg mg^−1^) as quantified by high-performance liquid chromatography (HPLC). This approach had low average prediction error (5.9%) across a gradient of astaxanthin contents and was only unreliable at very low contents (<0.6 μg mg^−1^). In addition, our machine learning model outperformed single or dual wavelength linear regression models even when the spectral data was obtained with a spectrophotometer coupled with an integrating sphere. Overall, this study proposes the use of HSI in combination with a CNN for precise non-invasive quantification of astaxanthin in cell suspensions.

## 1. Introduction

Microalgae are a diverse group of single-cell photosynthetic organisms that can serve as a biotechnological platform to produce high-value biomolecules and quality biomass. Their high photosynthetic efficiency and rapid growth, their ability to thrive in water that is unsuitable for human consumption and their high content of pigments, polyunsaturated fatty acids and other health promoting compounds have increased commercial interest in microalgae production over the past decades (Kuech et al. 2023). Currently, commercial production of microalgae in Europe focuses on high-value products for the supplement and nutraceutical industry (24%) as well as the cosmetics industry (24%) (Kuech et al. 2023). Common to these industries is the use of astaxanthin, a red-orange secondary ketocarotenoid that in microalgal cells serves as a scavenger of free radicals to protect against photooxidative stress (Li et al. 2023). The high antioxidant capacity of astaxanthin has been observed *in vivo* and *in vitro* (Liu et al., 2009; Xue et al., 2017), driving interest in its potential therapeutic applications for cardiovascular, neurodegenerative and immune diseases (Liu et al., 2023). Additionally, astaxanthin is used as a colorant and is responsible for the natural coloration observed in salmonids, shrimp, lobsters and crayfish (Lorenz and Cysewski, 2000). Microalgal production of astaxanthin is centred in the cultivation of the green algae *Haematococcus pluvialis* (Lorenz and Cysewski, 2000). Under stress conditions such as nutrient limitation and high light, this species can accumulate up to 7% of its dry weight as astaxanthin (Liyanaarachchi et al., 2020). Astaxanthin is not a single molecule since the presence of multiple conjugated double bonds allows for several possible geometric isomers (Subramanian et al. 2014) with the most abundant being all-E, followed by 9Z-, 13Z- and 15Z-forms in *H. pluvialis* (Subramanian et al. 2014; Koopmann et al., 2022). In microalgal cells, only small amounts of astaxanthin remain as free molecules (0.3-4%), while most of this ketocarotenoid is esterified with fatty acids to form monoesters (76– 94%) and diesters (2–23%) (Miao et al., 2006; Grewe and Griehl, 2008; Koopmann et al., 2022).

Monitoring of cultures is a critical aspect of microalgal production systems, since parameters such as biomass concentration and biochemical composition significantly influence the decision-making process of business operations (Solovchenko, 2023). Traditionally, microalgae culture monitoring depended on invasive methods in which a sample of biomass is taken, and different analytical laboratory methods are used to determine the parameter of interest. Currently however, efforts have been placed in developing non-invasive monitoring strategies to minimize the time lag between the measurement and availability of results as well as lowering the labour and operational costs required for the measurement (Havlik et al. 2022). Spectroscopic methods based on the absorption of light by microalgae remain the most widely used approach for routine monitoring of cultures (Li et al. 2020; Solovchenko, 2023). The Bouguer-Beer-Lambert law defines the linear correlation between the absorbance of a non-scattering medium and the sum of the products of each dissolved analyte’s molar absorptivity and its concentration, for a specified optical path length. This law is primarily applied to pigment extracts, but it can be an adequate approximation to study the absorbance of microalgal suspensions. Given that microalgal pigments absorb light in the visible range, the absorbance spectra between 350–700 nm can be used for the quantification of pigments while optical density (OD) in the near infrared (e.g OD_750_) can be used for whole biomass estimation (Griffiths et al. 2011). However, microalgal cell morphology and pigment packaging can affect light absorption and scattering by cell suspensions (Solovchenko et al. 2013), leading to non-linearities that deviate from the Bouguer-Beer-Lambert law. To minimize the impact of light scattering on a measured spectrum, an integrating sphere (IS) can be utilized, where most forward-scattered light from the sample is integrated into the measurement (Ritchie and Sma-Air, 2020). Nevertheless, a high-end IS can be expensive (Solovchenko, 2023).

Estimation of total astaxanthin content in *H. pluvialis* cultures is one of the crucial steps during commercial production. Commonly, cells are harvested, pigments extracted using physical disruption methods in combination with organic solvents and astaxanthin is subsequently quantified based on the light absorption properties of the total pigment extract (Boussiba and Vonshak, 1991). Pure all-E astaxanthin has a maximum absorption between 470–506nm depending on the solvent (Buchwald and Jencks, 1986), but there is significant overlap with the absorption peaks of chlorophylls and other carotenoids. This, added to the fact that stress conditions lead to the degradation of chlorophyll and the synthesis of secondary carotenoids other than astaxanthin (Ota et al., 2018), has prompted researchers to suggest using 530 nm as a wavelength to quantify astaxanthin while minimizing interference with the absorption peaks of other pigments (Casella et al., 2020). More sophisticated approaches for astaxanthin quantification require pigment extraction and de-esterification of astaxanthin followed by chromatographic separation and quantification of each astaxanthin isomer independently (Subramanian et al. 2014, Koopmann et al., 2022). Absorbance-based quantification is relatively fast, but due to the overlap of the astaxanthin absorption peak with other carotenoids, deviations of up to 30% in astaxanthin content can occur when astaxanthin content is lower than 2% of dry weight (Koopmann et al., 2022). This large error makes this estimation method very unreliable at low astaxanthin contents. In contrast, chromatography-based methods present high accuracy, but they are expensive, time consuming and require highly trained personal in addition to expensive equipment.

Reflectance hyperspectral imaging (HSI) has been proposed as a powerful and cost-effective alternative spectroscopic method for non-invasive monitoring of microalgal cultures (Murphy et al., 2013; Havlik et al. 2022; Salmi et al. 2022). A hyperspectral imager produces a set of images (>20) or data cube, where each image correspond to a different wavelength band and each pixel contains spatial information of the imaged target. One or two HSI wavelength bands can be used directly to, for example, quantify microalgal biomass (Murphy et al. 2014; Salmi et al. 2021; Pääkkönen et al. 2024). However, this approach misses the opportunity of utilizing the information contained across the whole visible spectrum. Machine learning (ML) algorithms represent an attractive approach to work with microalgal reflectance spectra since they can deal with the difficulties inherent to this kind of data such as the collinearity between close wavelengths. Additionally, ML can deal with non-linear relationships which might be more suitable for modelling reflectance data from scattering suspensions (Gardner, 2018). ML algorithms that have shown promising results when dealing with spectroscopic data are convolutional neural networks (CNN; Ghamisi et al., 2016: Malek et al., 2018; Salmi et al. 2022; Pääkkönen et al. 2024). CNN were originally developed for computer vision tasks and consists of one or several convolution layers followed by a deep-forward neural network (dense layers). The convolution kernels work as filters that select the most important features in the input while the dense layers perform the classification or regression task. CNN applied to microalgae reflectance HSI data have been successfully used for species classification (Salmi et al. 2022; Pääkkönen et al. 2024) as well as for quantification of biomass and pigments (Pyo et al. 2019; Salmi et al. 2022; Pääkkönen et al. 2024; Salmi et al. 2024).

In this study, we tested the use of a CNN model to predict the astaxanthin content per dry weight (μg mg^−1^) of *H. pluvialis* suspensions based on visual reflectance HSI data. As a reference, we tested the prediction error of linear models constructed using single or dual wavelengths from reflectance HSI or absorbance data from a high-end IS. We hypothesize that the variation in carotenoids and chlorophylls during stress conditions is coordinated in *H. pluvialis* and that such spectral information can be used to quantify astaxanthin accurately. In addition, we hypothesize that although a costly IS can produce high-quality absorbance spectra, reflectance HSI data in combination with the non-linear capabilities of a CNN can outperform linear regression models constructed using IS data.

## 2. Materials and Methods

### 2.1. Laboratory cultivations

The *H. pluvialis* strain used in this study was obtained from the Norwegian Culture Collection of Algae (NORCCA) and maintained as a stock monoculture in Modified Wright’s Cryptophyte (MWC) algae medium (Guillard and Lorenzen, 1972) with the addition of filter-sterilized (0.22 μm) vitamins (0.5 μg L^−1^ biotin [B7], 0.5 μg L^−1^ cyanocobalamin [B12], 0.5 μg L^−1^ pyridoxine [B6], 0.1 mg L^−1^ thiamine HCL [B1]). In the experimental set-up, cultivation was divided into two stages, with stage one prioritizing biomass accumulation while stress conditions were applied in stage two to maximize astaxanthin accumulation. In stage one, 8 parallel cultivations took place in 300 mL glass funnels aerated from the bottom (without additional CO_2_) as described in Stevčić *et al*. (2019) utilizing MWC media with higher concentrations of nitrogen and phosphorus (0.17 g L^−1^ of NaNO_3_ and 0.023 g L^−1^ of K_2_HPO_4_x3H_2_O). Light was supplied at 38–42 umol m^−2^ s^−1^ with LED growth lights (18 W, L-series T8 tubes, Valoya Oy) under a 12:12 photoperiod and room temperature was set at 20 °C. Cells were maintained in stage one until cultivations presented a green-brown colour. At that point, 200mL of culture were centrifuged for 5 min (2500 g, 18 °C) and cells were transferred to 500mL Erlenmeyer flasks containing 200mL of nitrogen deficient MWC media. Erlenmeyer flasks were placed in a shaker under constant high light (1400–1550umol m^−2^ s^−1^) supplied by LED lights (R150 LED Roof Lighting, Valoya Oy) and aerated with filtered room air. Fresh MWC media was added to the culture funnels (stage one) to replace the subtracted amount and such funnel was considered a new culture. Stage one lasted no more than two weeks from the addition of fresh media while stage two did not last more than three weeks. In total, 35 and 21 cultures were sampled from stage one and two, respectively.

### 2.2. Microalgal suspension spectroscopy

Sampling for spectroscopy was conducted using a pseudo-randomized approach, meaning that samples were not collected at consistent, predefined time intervals. Additionally, there was a focus on early stages of astaxanthin accumulation to obtain a dataset with spectral data of cell suspensions with low astaxanthin contents per dry weight. In practice, 100 mL of microalgae culture were collected at different timepoints from cultivation stages one and two to obtain its absorbance and reflectance spectrum. To increase the variability in astaxanthin concentration of the dataset, mixtures of samples from stages one and two were carried out by mixing different volumes of each sample (up to 1 and 99% of each culture). In total, 98 spectra were recorded of which 42 corresponded to sample mixtures, while the rest (56) corresponded to non-mixed samples (35 and 21 spectra from cultures of stage one and two, respectively).

Absorbance spectra were recorded with a spectrophotometer (Lambda 850 UV/VIS, Perkin Elmer) equipped with an integrating sphere by placing a glass cuvette (outer dimensions 50 × 50 × 14 mm, OP38, eCuvettes) containing the microalgal suspension in front of the integrating sphere. The cuvette had an optical path length of 10 mm (approximately 30 mL of microalgae). Spectra were measured across 350–750 nm with a wavelength resolution of 1 nm.

Reflectance spectra were recorded with a hyperspectral imager (Specim FX10, Specim, Finland) as described in Pääkkönen et al. (2024). Briefly, microalgal suspensions (40mL) were placed in 50mL cell culture flasks (EasYFlask, Nunc) at 25 cm from the camera lens. The camera and the light source (3 bulbs of DECOSTAR 51 ALU 20W 12V 36deg GU5.3 halogen) were mounted in a motorized scanner (LabScanner 40 × 20, Specim, Finland) and scanning speed was set to 2.0 mm s^−1^. White (PTFE diffuse reflector sheet PMR10P1, Thorlabs) and black references were imaged for each image separately. Spectra were measured across 400– 1000 nm with a spectral resolution of 5.5 nm

### 2.3. Biomass collection and astaxanthin quantification

After spectroscopical measurements, a known volume of microalgal suspension was filtered through a pre-weighed fibre filter (GF/A, Whatman) and subsequently re-weighed after 12 hr at 105 °C to determine its dry weight (mg L^−1^). The rest of the microalgal suspensions were transferred to 50 mL Falcon tubes and centrifuged for 10 min (5000 g, 18 °C). Cell pellets were transferred to 1.5 mL Eppendorf tubes and stored at –80 °C. Microalgal biomass was freeze-dried overnight before pigment extraction. Freeze-dried biomass (1 ± 0.1mg) was transferred to ZR BashingBead Lysis tubes (0.1 & 0.5 mm, Zymo research) and 3.1 μg of internal standard (Trans-β-Apo-8′-carotenal, Sigma Aldrich) were added together with 1 mL of ice-cold acetone. Tubes were homogenized for 20 seconds at 4 m s^−1^ for 4 rounds (Bead Ruptor Elite, OMNI) and subsequently centrifuged for 15 min (15000 RCF at 4 °C). The pigment-rich supernatant was placed in a 10 mL KIMAX tube and stored at –20 °C until further use. A second extraction round with ice-cold acetone, exactly as described, was carried out and supernatants were combined. To the final combined supernatants (approx. 2 mL), 1 mL of acetone, 2 mL of Tris solution 50 mM (pH = 7) and 0.6 mL of cholesterol esterase enzyme (3.3 U mL^−1^) isolated from *Pseudomonas* sp. (MP Biomedicals, Germany) were added. Freeze-dried enzyme was initially resuspended in Tris 50mM to achieve the desired concentration (3.3 U mL^−1^). Final enzyme concentration in KIMAX tube was 0.36 U mL^−1^ as suggested by Koopmann et al. (2022). De-esterification of astaxanthin was carried out at 37 °C in a water bath (in darkness) for 45 min. After de-esterification, samples were allowed to cool down to room temperature, 2 mL of petroleum ether were added and tubes were centrifuged for 3 min (2000 g) to facilitate phase separation. The pigment-rich upper phase was transferred to a new KIMAX tube and a second extraction with 2 mL of petroleum ether was carried out (as before). Upper phases were combined and evaporated under a nitrogen stream in total darkness. To dry tubes, 300 μl of acetone were added and pigment extracts were transferred to brown vials. During the whole extraction process, samples were protected from direct light exposure and only dim was used when necessary.

Pigment extracts were separated and analysed via high-performance liquid chromatography (HPLC; Nexera, Shimadzu) coupled with a SPD-M20A diode array detector (Shimadzu) using an Acclaim C30 column (3 μm 150 × 3.0 mm, Thermo Fisher Scientific). In total, three astaxanthin peaks were identified by their retention time and characteristic spectra (Subramanian et al. 2014) as all-E, 13Z-and 9Z-form (in decreasing order of abundance, respectively). It is likely that other less abundant astaxanthin isomers co-eluted with the three identified forms. Quantification of astaxanthin (μg) was done by interpolation in a 3-point calibration curve generated with synthetic (3S,3’S)-Astaxanthin standard (DHI). It was assumed that the area-to-concentration relationship of the astaxanthin standard was the same for all identified astaxanthin isomers. Astaxanthin (μg) was then normalized by the weighed dry weight to obtain astaxanthin content per dry weight (μg mg^−1^). To correct for the astaxanthin lost during the extraction process, the recovery percentage of the internal standard was used as a correction factor for astaxanthin. Sample recovery percentage was 77 % on average with two of the analysed samples been outliers (Fig. S1). These two outliers were not included in further analysis. Finally, astaxanthin concentration per litre was calculated by multiplying astaxanthin content in mg g^−1^ by the dry weight (g L^−1^) of the culture.

### 2.4. Linear regression

HSI data was first cropped to the region of interest (120×150 pixels) where the *H. pluvialis* suspension was located in each image (Fig. S2). Then, the dataset without outliers (n=96) was randomly divided into training and testing data (n = 76 and n = 20, respectively). Given that each ROI had 18,000 pixels, each with its own reflectance spectrum, the average reflectance was calculated for each training and test sample. Principal component analysis (PCA) of training samples was then carried out using the library sklearn and test samples were projected into the calculated principal components. The IS data partition was the same as with HSI data, meaning that the same samples were training and testing for both HSI and IS to ensure that the models created with both instruments were comparable. Selection of wavelengths for modelling was done using training data only. The absolute Pearson correlation was calculated between wavelengths from absorption or reflectance data with astaxanthin concentration per litre (mg L^−1^) and content per dry weight (μg mg^−1^). This was done by iterating over every single wavelength directly or a wavelength divided by 750nm and selecting the highest absolute correlation. A linear model was then built between the selected wavelength and astaxanthin in the corresponding unit using the Python library SciPy (module scipy.stats). The presence or absence of an intercept was decided based on which model presented the highest coefficient of determination (R^2^) with the training data. Model fitting on test data was evaluated with the root mean squared error (RMSE) and mean absolute percentage error (MAPE), calculated as follows:

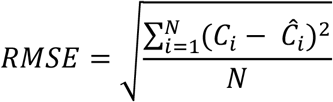

and

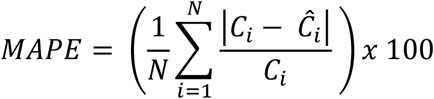

where *C*_*i*_ is the expected astaxanthin (μg L^−1^ or μg mg^−1^), Ĉ_*i*_is the predicted astaxanthin and *N* is the number of samples. To study if MAPE was affected by cultures with low astaxanthin, the MAPE of samples with quantified astaxanthin was >2 μg L^−1^ or μg mg^−1^ was calculated. This statistic is referred in the text as MAPE_>2_. Additionally, Pearson correlation and mean squared error (MSE) of test data was calculated.

### 2.5. Data augmentation and 1D Convolutional Neural Network

HSI training data (not averaged) was augmented by an iterative process where a random training sample was selected and the reflectance spectrum of 100 randomly selected pixels were averaged. This process was repeated 15000 times, to obtain 15000 augmented samples. To visualize the effect of data augmentation on the distribution on the training data, augmented data was projected into the principal components previously calculated (see “Linear regression” section). A portion of the augmented training data (20%) was randomly selected as validation data for model training. Only the wavelengths in the visible range were used for modelling (398.91 – 757.38nm) since the prediction error was lower than using the full HSI spectra (data not shown). Before model training, all spectral data (training, validation and test) were min-max-normalized.

A One-dimensional Convolutional Neural Network (1D CNN) was developed to predict astaxanthin content per dry weight (μg mg^−1^). The model was implemented with Python (v 3.10.9) using Keras library (v 3.1.1) and Tensorflow (2.16.1) backend on Jupyter lab (3.5.3) editing environment. A simplified custom CNN architecture was employed, which omitted the use of max-pooling layers (Springenberg et al., 2014) and connected each neuron in the last convolution layer to only one neuron in the first dense layer (Table S1, Fig. 1). By using this architecture, the number of trainable parameters was reduced from 208,881 to 31,051. Convolution layers had no activation function while dense layers, except for the output layer, used Leaky Rectified Linear Unit (Leaky ReLU). The output layer had a linear activation function. To avoid overfitting, each neuron in the dense layers had a dropout rate of 10%. The model was trained with a gradient-based stochastic optimizer (Adam) using a learning rate of 0.003 for a maximum of 200 epochs (with early stopping). Batch size was 30 samples and mean squared error (MSE) was used as a loss function. The number of convolution and dense layers were manually adjusted until high performance was achieved. Once the best model was obtained (based on lowest MSE of validation data), the SHAP algorithm (SHapley Additive exPlanations) was used to analyse which areas of the spectra were important for the model prediction.

**Fig. 1.**
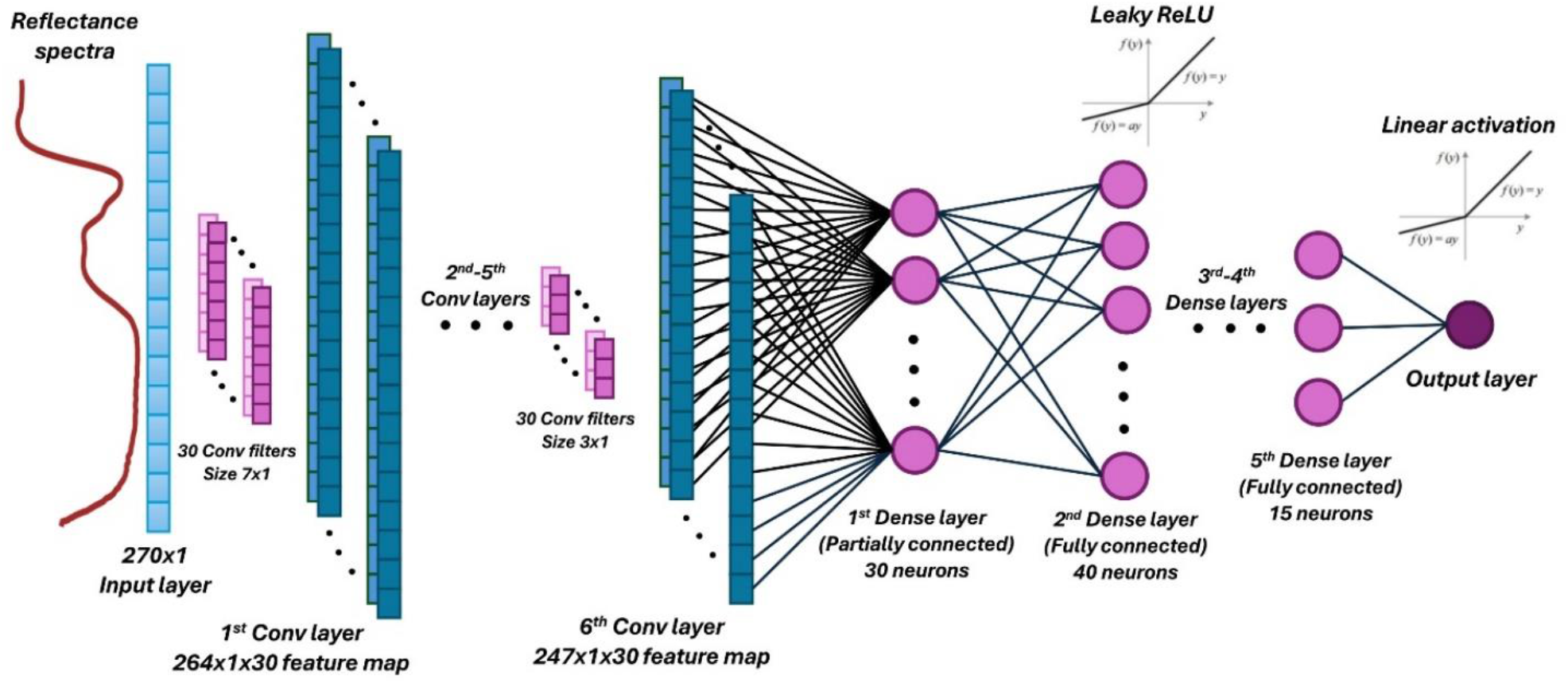
1-Dimensional (1D) convolutional neural network (CNN) architecture to predict astaxanthin content per dry weight (μg mg^−1^) utilizing reflectance hyperspectral imaging (HSI) data of *H. pluvialis* suspensions.

## 3. Results

### 3.1. Dataset characteristics and recorded spectra

Dry weight of *H. pluvialis* cultures ranged between 0.13-1.77 g L^−1^ with most samples having a dry weight <0.8 g L^−1^ (Fig. S3a). Total astaxanthin content ranged from 0.01-1% of dry biomass, with roughly half of the samples in the dataset presenting contents in the range of 0.01-0.2% (Fig. S3b). Of the three identified and quantified astaxanthin isomers, all-E astaxanthin constituted on average 86% of the total astaxanthin content, with approximately 10% attributed to 13Z-form and ∼4% of 9Z-form (Fig. S3c). Both IS and HSI spectra presented clear differences between *H. pluvialis* cultures with increasing stress conditions (Fig. 2). Although the use of an IS allowed to obtain high spectral resolution and an overall high quality absorbance spectrum, scattering of light markedly increased in stressed cells (Fig. 2a). This led to shifts in peak maxima which was not observed in HSI spectra (Fig. S4).

**Fig. 2.**
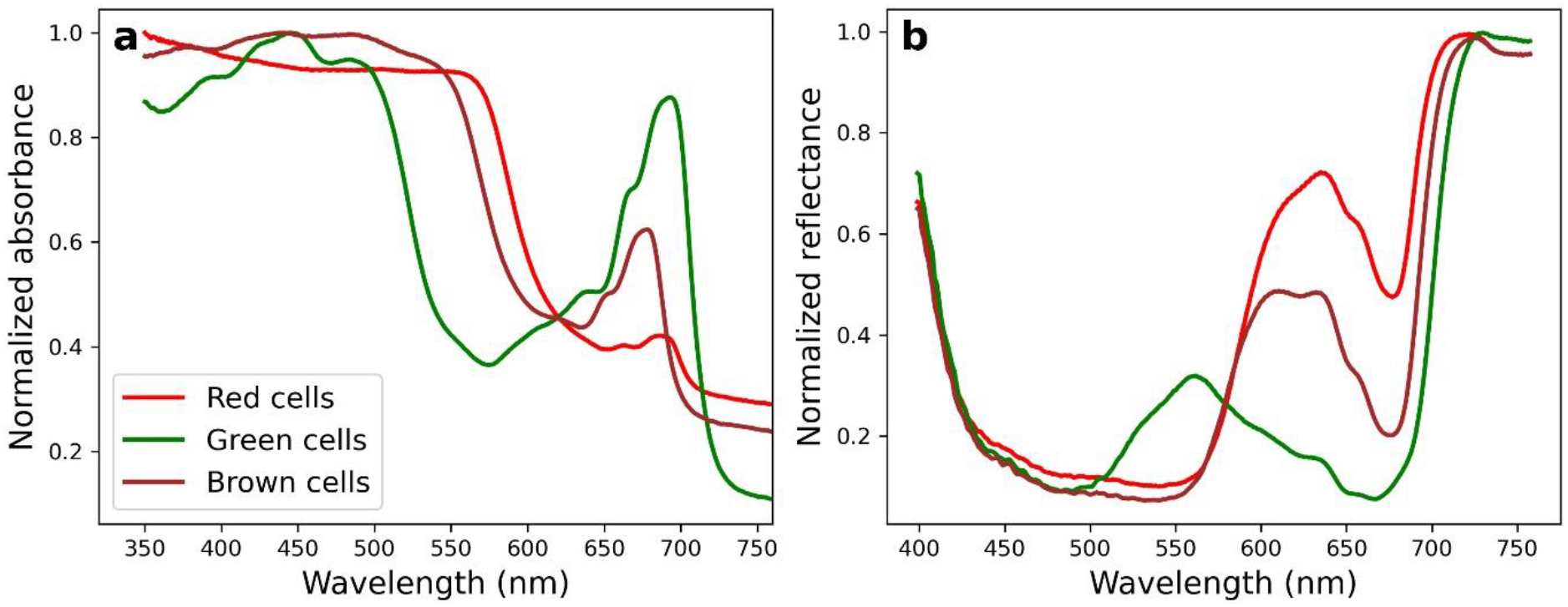
Examples of normalized absorption (**a**) and reflectance (**b**) spectrum of *H. pluvialis* suspensions at different cultivation times. With longer cultivation time and stress conditions, cultivations transition from green to brown and then to red colour. Reflectance data was acquired with a hyperspectral imager while absorbance data was obtained with a spectrophotometer attached to an integrating sphere.

### 3.2. Single and dual wavelength regression

Astaxanthin concentration per litre (mg L^−1^) reflects the total volumetric amount of astaxanthin and should therefore best correlate with a single wavelength. The highest absolute Pearson correlations were observed with wavelengths close to 565 nm for both IS (568 nm, r=0.91; Table S2) and HSI (564 nm, r=0.90; Table S2). Linear models using these highly correlating wavelengths did not accurately predict test samples and had MAPE of 432% and 214.3% for IS and HSI, respectively (Fig. 3A,B; Table S3; Table S4). Since these linear models showed poor fitting at low astaxanthin (<2 mg L^−1^), MAPE was calculated for test samples with higher concentrations (>2 mg L^−1^; n=8). MAPE_>2_ was notably lower for IS (27.4%) and HSI (35.3%), reflecting lower prediction error at higher astaxanthin concentrations (Table S4). However, astaxanthin concentration per litre might not be an ideal unit for culture monitoring given the effect of cell density. Therefore, an ideal model would predict the astaxanthin content per dry weight (μg mg^−1^) directly. Absolute Pearson correlation between a single wavelength and astaxanthin content per dry weight was highest with 560nm for both IS and HSI (r=0.67 and 0.88, respectively; Table S2). Linear models of 560nm and astaxanthin (μg mg^−1^) showed a lower MAPE than previous models (145.4% and 76% for IS and HSI, respectively; Figure 3C,D; Table S3; Table S4), although their performances at >2 μg mg^−^1 were similar (MAPE_>2_ = 41.1% and 36.6% for IS and HSI, respectively). Lastly, the use of two wavelengths was tested using optical density at 750nm (OD 750) as a normalizing value given its high correlation with dry weight (r=0.99 for both IS and HSI; Table S3, Fig. S5). The highest Pearson correlations using this approach were observed at 497/750 for IS and 560/750 for HSI (r= 0.84 and 0.83 for IS and HSI, respectively; Table S2). Linear models using these dual wavelengths gave >100% MAPE (116.3% for IS and 120.8% for HSI; Table S4) and similar results as the previous tested models at higher astaxanthin contents (MAPE_>2_= 24.7% and 35% for IS and HSI, respectively; Figure 3; Table S3; Table S4).

**Fig. 3.**
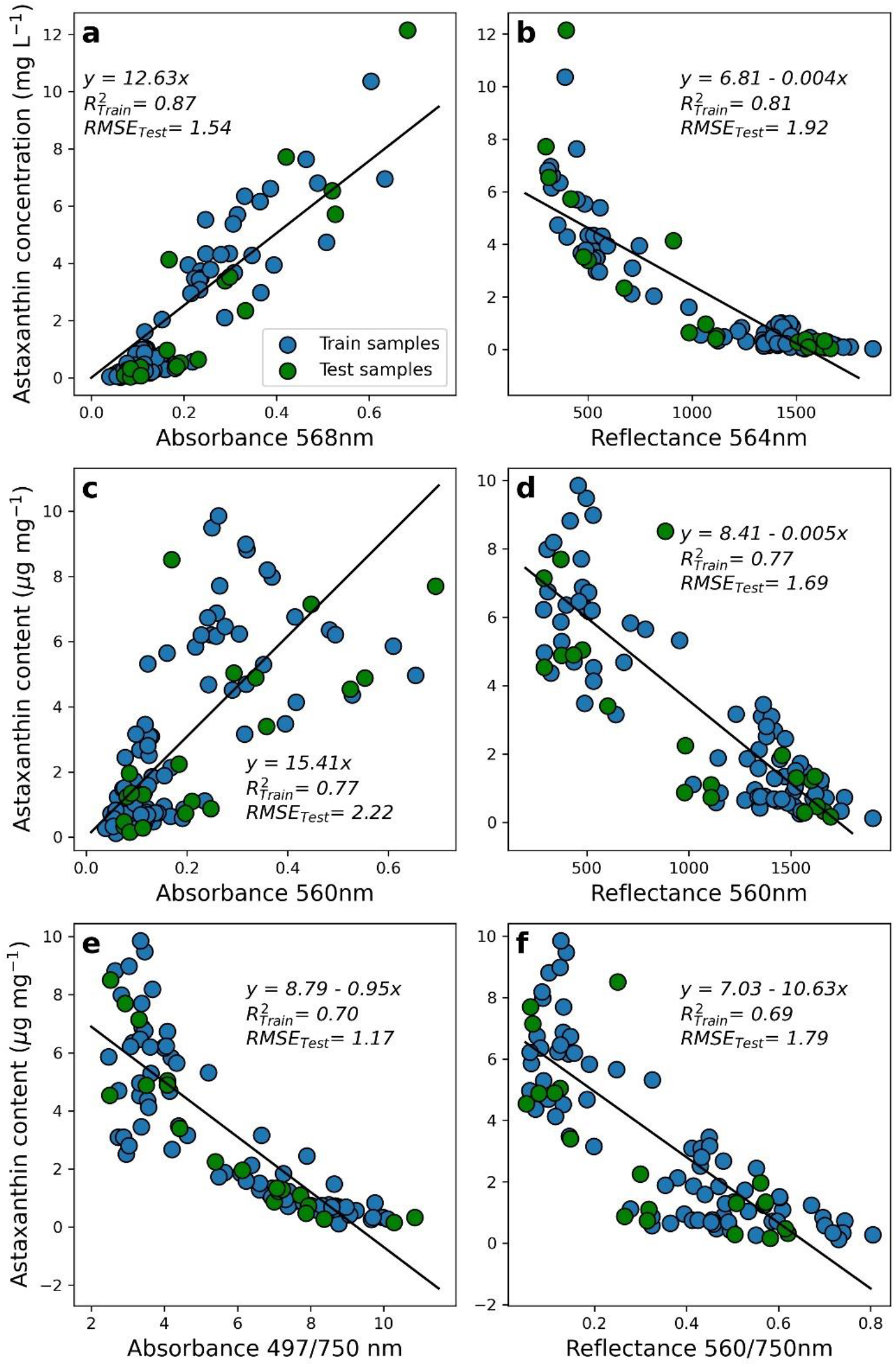
Linear models describing the relationship between single (**a**-**d**) or dual (**e**,**f**) wavelengths and astaxanthin concentration per litre (**a**,**b**) or content per dry weight (**d**-**f**). Reflectance data was acquired with a hyperspectral imager while absorbance data was obtained with a spectrophotometer attached to an integrating sphere. Linear equation, train data coefficient of determination (R^2^) and test data root mean squared error (RMSE) are presented in each plot.

### 3.3. 1D Convolutional neural network regression

The use of a CNN for the prediction of astaxanthin content per dry weight (μg mg^−1^) using HSI data strongly decreased the prediction error compared to single and dual wavelength linear model (Figure 4). Pearson correlation between expected and predicted astaxanthin content was r=0.99 (Table S3). The CNN had the lowest MAPE of all studied models in this work (26.8%) and presented a very low prediction error for astaxanthin content >2 μg mg^−1^ (MAPE_>2_ = 5.9%; Table S4). On average, the test samples with <0.6 μg mg^−1^ astaxanthin (or 0.06% of biomass) had >10 times prediction error than samples with >0.6 μg mg^−1^ astaxanthin (Fig. 4b), suggesting that at such low astaxanthin contents accurate predictions might not be possible using HSI. PCA summarized 95% of the variance observed between sample’s spectra in the first two principal components (Fig. 5A). Position of train and test samples in the two principal component axes shows that train samples represent well the variability observed in test samples, hence suggesting a good generalization capability of the CNN model. Plotting of the augmented spectra used for model training shows a larger portion of the two-dimensional space covered (Fig. 5B), validating the use of random pixel selection to increase the variability in training data during the model training process. Shapley-based explanation of the CNN model shows that the most important wavelengths used for model prediction are in the range of 550-700nm (Figure 5C), with 578.28, 584.99 and 630.79nm presenting the highest contribution to the prediction value. A closer inspection at the effect of the 20 most important features for value prediction suggest that when wavelengths >580nm have low values they have a negative effect on the astaxanthin content while at high values they have a positive effect on the prediction value (Fig. 5D). Since the second absorption peak of chlorophylls are in this area of the spectrum, it can be interpreted that chlorophyll concentration is negatively related to the predicted content of astaxanthin. The opposite trend is seen for wavelengths <580nm, suggesting that lower values in the carotenoid area of the spectra increase the predicted astaxanthin content by the CNN while higher values decrease the prediction estimate.

**Fig. 4.**
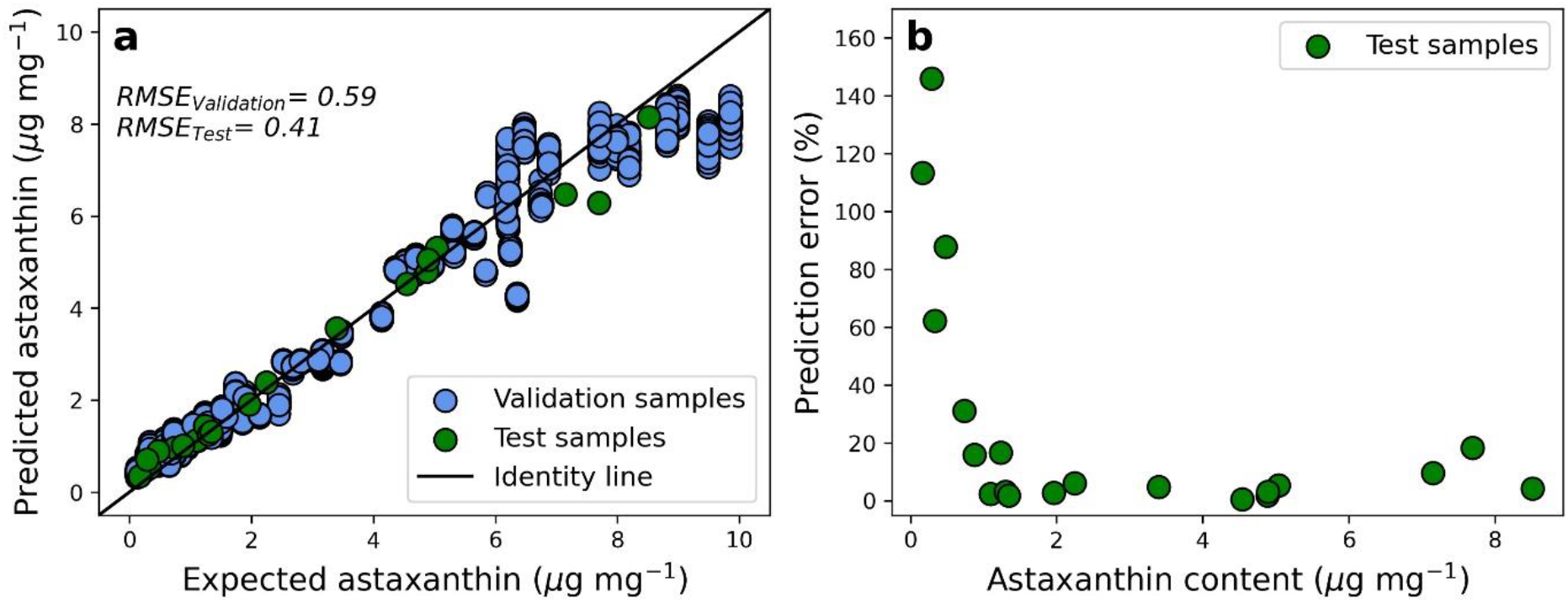
Relationship between the predicted astaxanthin content (μg mg^−1^) using a convolutional neural network (CNN) and the real values of validation and test datasets (**a**) together with the dependency of absolute prediction error with the astaxanthin concentration of test samples (**b**).

**Fig. 5.**
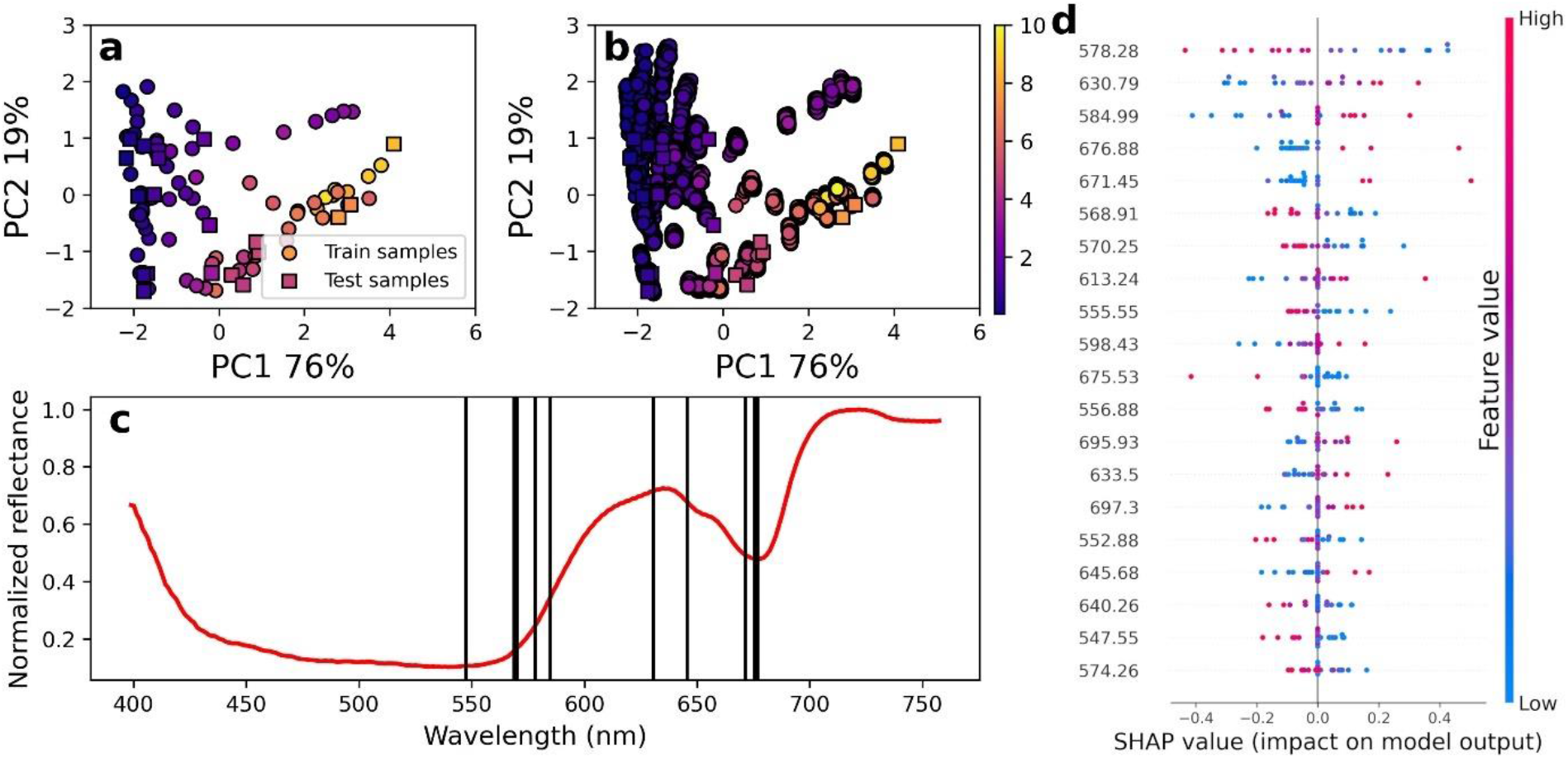
Principal component analysis (PCA) of train and test samples without (**a**) or with augmented training data (**b**). Variance explained by each principal component is given as percentage in plot axes and colour coding reflects the astaxanthin content (μg mg^−1^) of each sample as indicated in the colour bar (**a, b**). Most influential wavelengths (10) for prediction of astaxanthin identified by SHAP algorithm are plotted as vertical black lines on top of a normalized reflectance spectrum (red) of *H. pluvialis* suspension (**c**). Impact of wavelength value on model prediction value (**d**) of the 20 most influential wavelengths (ordered from higher to lower influence).

## 4. Discussion

Astaxanthin quantification from *H. pluvialis* cultures requires laborious sample processing while accuracy at low astaxanthin contents can only be achieved using HPLC (Koopmann et al., 2022). This study shows that applying machine learning algorithms to predict astaxanthin content using reflectance HSI of *H. pluvialis* suspensions is an effective approach for fast and accurate non-invasive quantification. Traditionally, spectroscopic analysis of microalgal suspensions has focussed on biomass monitoring using absorbance data (i.e., OD750) in combination with appropriate calibration curves (Griffiths et al. 2011). Nevertheless, pigment-specific quantification is more challenging due to partial or total overlap of pigment absorption peaks, scattering of light by microalgal cells and shifts in pigment absorption maximum due to intermolecular interactions (Solovchenko et al., 2013). Our reflectance HSI data showed no major optical effects due to changes in cell density or culture conditions, although the chlorophyll absorption peak in the red region exhibited a slight rightward shift in stressed cells. In contrast, IS spectra presented clear effects of light scattering in stressed cells. Nitrogen deficiency and high light, common conditions used to stimulate astaxanthin production, promote the encystment of *H. pluvialis* cells (Kobayashi et al. 1997; Ota et al., 2018) leading to changes in cell size, chlorophyll degradation and the accumulation of secondary carotenoids in lipid droplets (Ota et al., 2018). Such morphological changes have been shown to increase the scattering coefficient of *H. pluvialis* suspensions (Sheng et al. 2018). Despite an IS technically captures all forward-scattered light from a sample, our results show that the contribution of side-and back-scattering of red *H. pluvialis* suspensions can affect the quality of the obtained spectra leading to the overestimation of light absorption and shifts in peak absorption maxima.

The astaxanthin absorption peak overlaps to a high degree with other carotenoids present in *H. pluvialis* such as lutein and β-carotene (Casella et al. 2020). Therefore, the use of higher wavelengths (i.e. 530 nm) than the absorbance maximum of astaxanthin have been suggested for the quantification of pigment extracts from *H. pluvialis* (Casella et al. 2020). Our Pearson correlation analysis between astaxanthin concentration per litre (mg L^−1^) or content per dry weight (μg mg^−1^) with a single wavelength favoured the use of ≥560nm. These results suggest that the range between 550-600nm contains wavelengths where astaxanthin absorption has less overlap with other pigments i*n vivo*. Nevertheless, precise determination of the astaxanthin absorption peak *in vivo* is difficult due to the presence of different astaxanthin esters (Grewe and Griehl, 2008) and the possibility of spectral changes due to strong pigment aggregation in stressed cells (Solovchenko et al., 2013). Prediction errors of linear models using single or dual wavelengths were high although large improvements were seen at higher astaxanthin amounts (>2 mg L^−1^ or >2 μg mg^−1^). This difference in prediction error was expected due to the very low astaxanthin amounts observed in some of the samples analysed. Considering that reflectance and absorption of a highly scattering suspension are not linearly related to changes in the concentration of the light absorbing analyte (Gardner, 2018), it is likely that non-linear models would increase the prediction accuracy of the tested single and dual wavelengths. Nevertheless, since *H. pluvialis* presents different scattering coefficients at different life stages (Sheng et al. 2018) and the overlap between the astaxanthin absorption peak and other carotenoids is so pronounced, it is possible that no single or dual wavelength model could produce very accurate astaxanthin estimations even when using a high-end IS.

Although CNNs were originally developed as an image classification algorithm (LeCun et al., 1998), their suitability to deal with 1-dimensional spectroscopic data has also been demonstrated (Malek et al., 2018; Salmi et al. 2022; Pääkkönen et al. 2024). Our results provide further evidence that CNNs can handle the collinearity of spectroscopic data and produce accurate regression results. Estimation of astaxanthin content per dry weight had an average prediction error 3-fold lower than the best tested linear model using either HSI or IS data. Moreover, our CNN model outperformed the accuracy of spectrometric quantification of astaxanthin using pigment extracts at low astaxanthin contents (Koopmann et al., 2022). This improvement in prediction error is likely explained by the capacity of the CNN to extract the important features from the HSI data and model a non-linear relationship between such features and the output. By doing so, the CNN model decreases its reliance on a single wavelength such as an absorption maximum which, as previously mentioned, can be subjected to spectral shifts due to changes in the spectral properties of the algal culture. The non-linearity of the HSI data is evidenced by our PCA results, where astaxanthin content presents a non-linear distribution in principal component axes. As observed in the SHAP values, the CNN is not only using the wavelengths associated with astaxanthin but also the red absorption of chlorophylls to make predictions. The relationship between total carotenoids and chlorophyll has been shown to relate non-linearly to OD_500_/OD_678_ (Solovchenko et al., 2013), suggesting that the CNN model leverages such relationship to estimate astaxanthin content.

Common downsides to the use of CNNs are the large amounts of data and computing power required for model training. In this study, we overcame these limitations by augmenting our training dataset and utilizing a custom CNN architecture that significantly decreases the number of trainable parameters. Overall, our data augmentation strategy combined with the use of random dropout of neurons during training resulted in good generalization of the CNN model despite the low number (n=76) of independent training samples. Limitations to our study include the ranges of cell density and astaxanthin content of our samples. Unfortunately, higher biomasses or astaxanthin content per dry weight could not obtained under our laboratory conditions. More research on the ability of HSI and CNN to predict astaxanthin content under relevant commercial conditions is needed.

## 5. Conclusions

Reflectance HSI in combination with CNN offers a fast and cost-effective non-invasive monitoring approach to astaxanthin content quantification of *H. pluvialis* suspensions. The low prediction error observed in this study was unmatched by simple linear models utilizing one or two wavelength bands of the spectral data, even when using a high-end integrating sphere for absorbance spectrum acquisition. Data augmentation and a simplified CNN architecture allowed to obtain good model generalization capabilities without the need for large amounts of training data or computing power. Future research on the detection limits of HSI of *H. pluvialis* commercial cultures will be key to define if the approach used in this study can translate to the industrial scale production of astaxanthin.

## CRediT author statement

**Marco L. Calderini:** Conceptualization, Methodology, Investigation, Software, Writing - Original Draft, Review & Editing. **Salli Pääkkönen:** Investigation, Review & Editing. **Aliisa Yli-Tuomola:** Investigation, Review & Editing. **Hemanta Timilsina:** Methodology, Review & Editing. **Katja Pulkkinen:** Resources, Review & Editing. **lkka Pölönen:** Resources, Review & Editing. **Pauliina Salmi:** Conceptualization, Supervision, Funding acquisition, Review & Editing.

## Acknowledgements

The authors would like to thank laboratory technicians Mervi Koistinen and Emma Pajunen and laboratory engineer Hannu Pakkanen for their help during the experimental work.

## Appendix A. Supplementary data

The following file contains the Supplementary information to this article.

## Data availability

All data required to reproduce the results and figures of this manuscript will be publicly available in the IDA database (Fairdata.fi) upon publication.

