## Supplemental file (figures and tables) for "Accurate non-invasive quantification of astaxanthin content using hyperspectral images and machine learning"

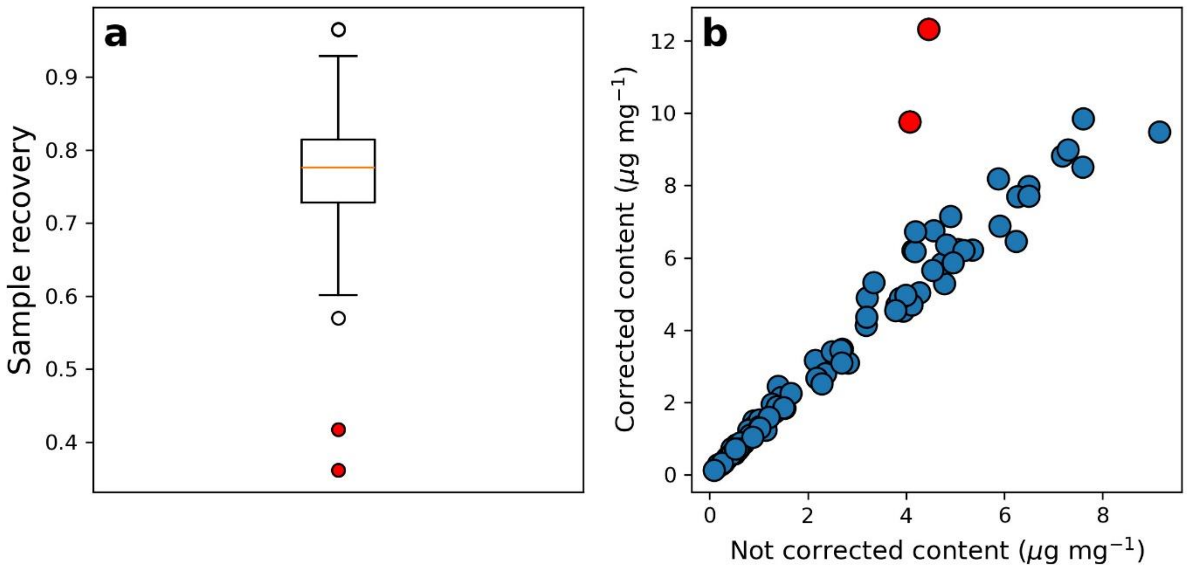


**Fig. S1** Astaxanthin recovery percentage (**a**) and correction of astaxanthin concentent based on sample recovery percentage (**b**). The box in the presented boxplot shows the quartiles of the dataset and the whiskers extend to show the rest of the distribution while the median value is presented in orange. Outliers are presented as open circles and extreme outliers are coloured red.

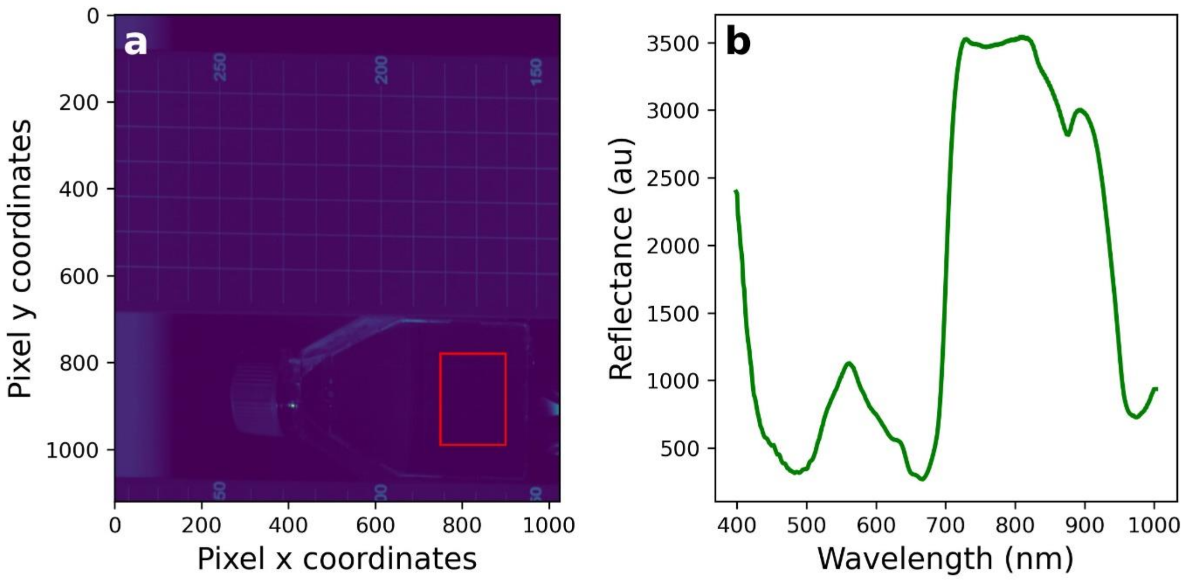


**Fig. S2** Example of hyperspectral image (**a**) and the selected region of interest (ROI; red square) together with the average reflectance spectra of all pixels in the ROI (**b**).

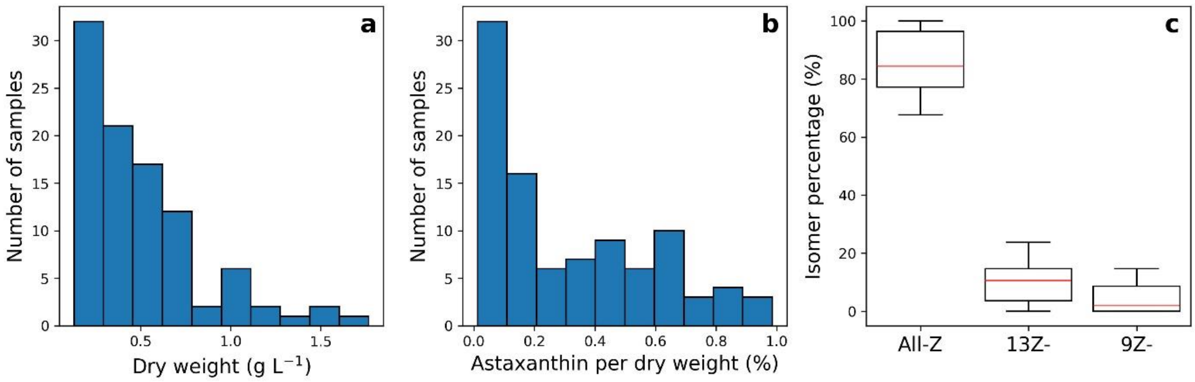


**Fig. S3** Distribution of dry weight (g L$^{-1}$; **a**) and total astaxanthin content as percentage of dry weight (**b**) in the dataset. The contribution of each astaxanthin isomer (**c**) to the total astaxanthin is presented as a boxplot where the median values are presented in orange, the box shows the quartiles of the dataset and the whiskers extend to show the rest of the distribution.

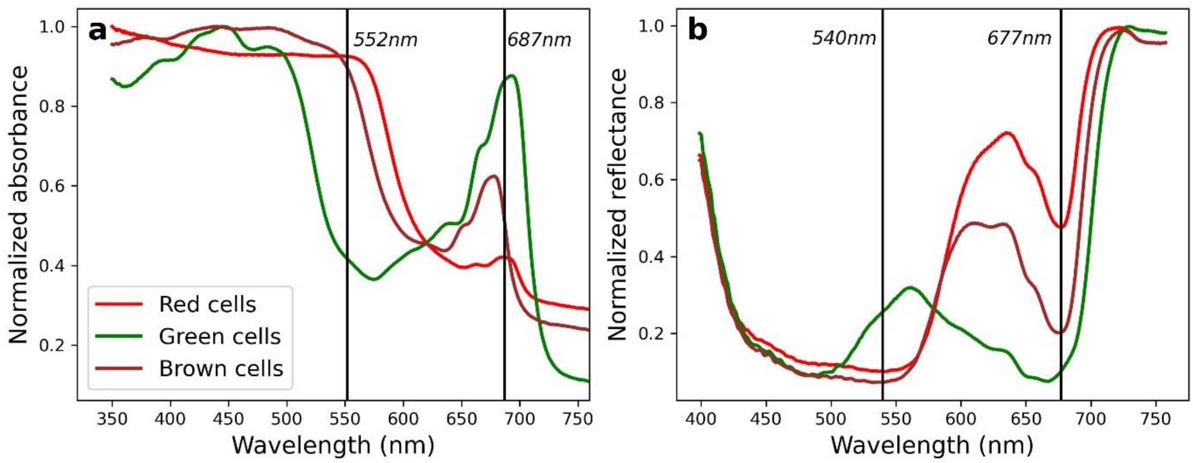


**Fig. S4** Shifts in position of peak maxima for carotenoids and red-absorption of chlorophyll in absorbance spectra obtained with an integrating sphere (**a**) and reflectance spectra obtained with a hyperspectral imager (**b**). Vertical lines correspond to the wavelength where the maximum absorbance of the carotenoid peak and the red peak of chlorophyll are maximum. Vertical lines were calculated using the spectrum of red cells (red colour spectrum). Legend is shared between plots.

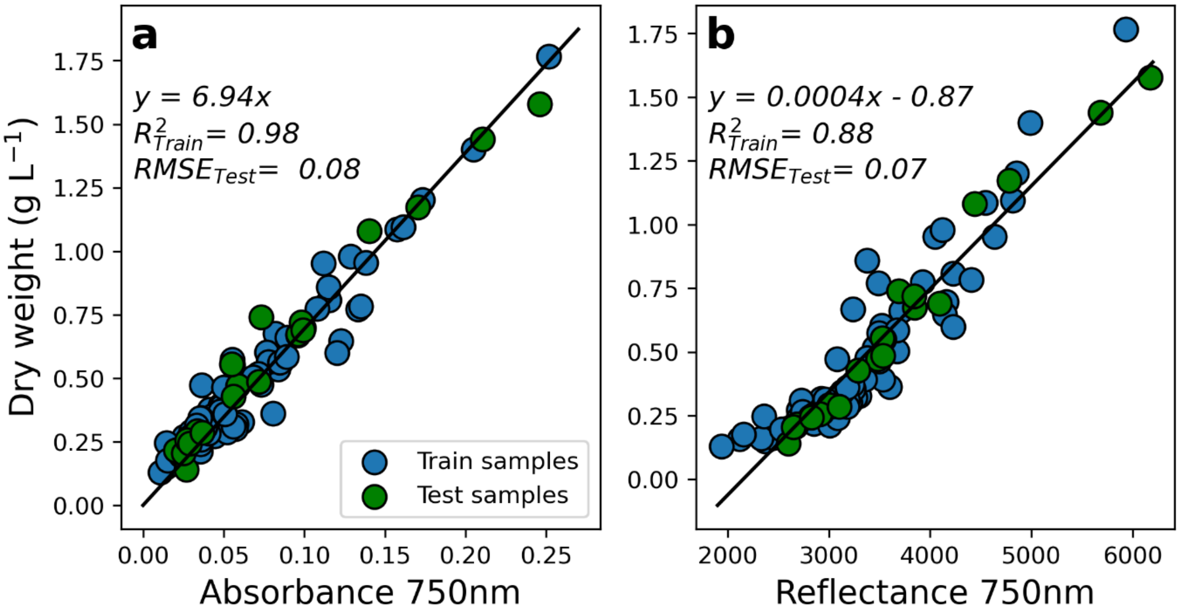


**Fig. S5** Relationship between absorbance (**a**) or reflectance (**b**) at 750nm and dry weight (g $L^{-1}$). Absorbance data was obtained spectrophotometrically using an integrating sphere while reflectance data was obtained using a hyperspectral imager. Legend is shared between plots.

**Table S1** Detailed 1-Dimensional (1D) convolutional neural network (CNN) architecture to predict astaxanthin content per dry weight (μg mg−1) utilizing reflectance hyperspectral imaging (HSI) data of H. pluvialis suspensions.

| Layer | Kernel size | Activation | Output shape | Parameters |
|---|---|---|---|---|
| Conv1D | 7 | None | (None,264,30) | 240 |
| Conv1D | 5 | None | (None,260,30) | 4530 |
| Conv1D | 5 | None | (None,256,30) | 4530 |
| Conv1D | 5 | None | (None,252,30) | 4530 |
| Conv1D | 4 | None | (None,249,30) | 3630 |
| Conv1D | 3 | None | (None,247,30) | 2730 |
| Flatten | | | (None,7410) | 0 |
| Dense | | LeakyReLU | (None,1) | 248 |
| Dense | | LeakyReLU | (None,1) | 248 |
| Dense | | LeakyReLU | (None,1) | 248 |
| Dense | | LeakyReLU | (None,1) | 248 |
| Dense | | LeakyReLU | (None,1) | 248 |
| Dense | | LeakyReLU | (None,1) | 248 |
| Dense | | LeakyReLU | (None,1) | 248 |
| Dense | | LeakyReLU | (None,1) | 248 |
| Dense | | LeakyReLU | (None,1) | 248 |
| Dense | | LeakyReLU | (None,1) | 248 |
| Dense | | LeakyReLU | (None,1) | 248 |
| Dense | | LeakyReLU | (None,1) | 248 |
| Dense | | LeakyReLU | (None,1) | 248 |
| Dense | | LeakyReLU | (None,1) | 248 |
| Dense | | LeakyReLU | (None,1) | 248 |
| Dense | | LeakyReLU | (None,1) | 248 |
| Dense | | LeakyReLU | (None,1) | 248 |
| Dense | | LeakyReLU | (None,1) | 248 |
| Dense | | LeakyReLU | (None,1) | 248 |
| Dense | | LeakyReLU | (None,1) | 248 |
| Dense | | LeakyReLU | (None,1) | 248 |
| Dense | | LeakyReLU | (None,1) | 248 |
| Dense | | LeakyReLU | (None,1) | 248 |
| Dense | | LeakyReLU | (None,1) | 248 |
| Dense | | LeakyReLU | (None,1) | 248 |
| Dense | | LeakyReLU | (None,1) | 248 |
| Dense | | LeakyReLU | (None,1) | 248 |
| Dense | | LeakyReLU | (None,1) | 248 |
| Dense | | LeakyReLU | (None,1) | 248 |
| Dense | | LeakyReLU | (None,1) | 248 |
| Concatenate | | LeakyReLU | (None,30) | 0 |
| Dropout (0.1) | | LeakyReLU | (None,30) | 0 |
| Dense | | LeakyReLU | (None,40) | 1240 |
| Dropout (0.1) | | LeakyReLU | (None,40) | 0 |
| Dense | | LeakyReLU | (None,30) | 1230 |
| Dropout (0.1) | | LeakyReLU | (None,30) | 0 |
| Dense | | LeakyReLU | (None,20) | 620 |
| Dropout (0.1) | | LeakyReLU | (None,20) | 0 |
| Dense | | LeakyReLU | (None,15) | 315 |
| Dropout (0.1) | | LeakyReLU | (None,15) | 0 |
| Dense | | Linear | (None,1) | 16 |

**Table S2** Highest absolute Pearson correlation (r) between a single or dual wavelength with astaxanthin concentration per litre (mg $L^{-1}$) or content per dry weight (μg $mg^{-1}$). Results are presented for reflectance hyperspectral imaging (HSI) and absorbance integrating sphere (IS) data. All correlations were calculated using only training data. In bold p-values lower than 0.01.

| Instrument | Astaxanthin unit | Wavelength (nm) | r | p-value |
|---|---|---|---|---|
| HSI | mg $L^{-1}$ | 563.56 | 0.899 | **<0.01** |
| | μg $mg^{-1}$ | 559.55 | 0.878 | **<0.01** |
| | μg $mg^{-1}$ | 560.89/750 | 0.829 | **<0.01** |
| IS | mg $L^{-1}$ | 568 | 0.914 | **<0.01** |
| | μg $mg^{-1}$ | 560 | 0.670 | **<0.01** |
| | μg $mg^{-1}$ | 497/750 | 0.837 | **<0.01** |

**Table S3** Statistics of linear models between a single or dual wavelength and biomass per litre (g $L^{-1}$), astaxanthin concentration per litre (mg $L^{-1}$), or content per dry weight (µg $mg^{-1}$) together with convolutional neural network (CNN). Pearson correlation (r), coefficient of determination ($R^2$) and root mean squared error (RMSE) are presented, clarifying if training, validation or testing data was used. Results are presented for reflectance hyperspectral imaging (HSI) and absorbance integrating sphere (IS) data. In bold p-values lower than 0.01.

| Instrument | Model | Statistic | Value | p-value |
|---|---|---|---|---|
| IS | Abs 750 – dry weight (g $L^{-1}$) | Intercept | - | - |
| | | Slope | 6.938 | **<0.01** |
| | | $R^2$ - Train | 0.977 | - |
| | | RMSE - Test | 0.084 | - |
| | | r - Test | 0.987 | **<0.01** |
| | Abs 560 – astaxanthin (µg $mg^{-1}$) | Intercept | - | - |
| | | Slope | 15.407 | **<0.01** |
| | | $R^2$ - Train | 0.765 | - |
| | | RMSE - Test | 2.220 | - |
| | | r - Test | 0.716 | **<0.01** |
| | Abs 568 – astaxanthin (mg $L^{-1}$) | Intercept | - | - |
| | | Slope | 12.631 | **<0.01** |
| | | $R^2$ - Train | 0.867 | - |
| | | RMSE - Test | 1.536 | - |
| | | r - Test | 0.924 | **<0.01** |
| | Abs 497/750 – astaxanthin (mg $L^{-1}$) | Intercept | 8.796 | **<0.01** |
| | | Slope | -0.949 | **<0.01** |
| | | $R^2$ - Train | 0.701 | - |
| | | RMSE - Test | 1.166 | - |
| | | r - Test | -0.897 | **<0.01** |
| HSI | 750nm – dry weight (g $L^{-1}$) | Intercept | -0.870 | **<0.01** |
| | | Slope | 0.000 | **<0.01** |
| | | $R^2$ - Train | 0.859 | - |
| | | RMSE - Test | 0.067 | - |
| | | r - Test | 0.988 | **<0.01** |
| | Wl 560 – astaxanthin (µg $mg^{-1}$) | Intercept | 8.409 | **<0.01** |
| | | Slope | -0.005 | **<0.01** |
| | | $R^2$ - Train | 0.770 | - |
| | | RMSE - Test | 1.693 | - |
| | | r - Test | -0.801 | **<0.01** |
| | Wl 564 – astaxanthin (mg $L^{-1}$) | Intercept | 6.811 | **<0.01** |
| | | Slope | -0.004 | **<0.01** |
| | | $R^2$ - Train | 0.808 | - |
| | | RMSE - Test | 1.925 | - |
| | | r - Test | -0.818 | **<0.01** |
| | Wl 560/750 – astaxanthin (µg $mg^{-1}$) | Intercept | 7.079 | **<0.01** |
| | | Slope | -10.697 | **<0.01** |
| | | $R^2$ - Train | 0.687 | - |
| | | RMSE - Test | 1.786 | - |
| | | r - Test | -0.772 | **<0.01** |
| | CNN – astaxanthin (µg $mg^{-1}$) | RMSE - Test | 0.405 | - |
| | | RMSE - Validation | 0.593 | **-** |
| | | r - Test | 0.993 | **<0.01** |

**Table S4** Mean absolute percentage error (MAPE) results of linear models and convolutional neural network (CNN). Results are separated for reflectance hyperspectral imaging (HSI) and absorbance integrating sphere (IS) data. MAPE calculated using samples with concentrations >2 (mg $L^{-1}$ or µg $mg^{-1}$) is presented as $MAPE_{>2}$.

| Instrument | Model | MAPE | $MAPE_{>2}$ |
|---|---|---|---|
| IS | 568nm – astaxanthin (mg $L^{-1}$) | 431.9 | 27.41 |
| IS | 560 – astaxanthin (µg $mg^{-1}$) | 145.4 | 41.07 |
| IS | 497/750 – astaxanthin (µg $mg^{-1}$) | 116.3 | 24.73 |
| IS | 560/690nm – astaxanthin (µg $mg^{-1}$) | 103.7 | 22.07 |
| HSI | 564nm – astaxanthin (mg $L^{-1}$) | 214.3 | 35.30 |
| HSI | 560 – astaxanthin (µg $mg^{-1}$) | 76.0 | 36.62 |
| HSI | 530/750 – astaxanthin (µg $mg^{-1}$) | 120.8 | 35.01 |
| HSI | 561/677nm – astaxanthin (µg $mg^{-1}$) | 79.6 | 23.27 |
| HSI | CNN – astaxanthin (µg $mg^{-1}$) | 26.8 | 5.97 |
